# Early synaptic pathology is associated with small tau aggregates in Alzheimer’s disease

**DOI:** 10.1101/2025.09.13.676039

**Authors:** Emre Fertan, Shekhar Kedia, Georg Meisl, Matthew W. Cotton, Annelies Quaegebeur, Maria Grazia Spillantini, David Klenerman

**Author notes:** Correspondence to: Professor Sir David Klenerman, Yusuf Hamied Department of Chemistry, University of Cambridge, UK, CB2 1EW. Equally contributed co-first authors (Emre Fertan, Shekhar Kedia).

## Abstract

Alzheimer’s disease (AD) is phenotypically characterised by progressive memory loss, which involves tau aggregation and synaptic pathology. Here we characterised the tau aggregates in individual synaptosomes from AD cases and controls measuring their number and size using single molecule fluorescence microscopy. A total of 7,888 synaptosomes from pre-frontal cortex samples were studied, showing the presence of AT8-positive tau aggregates in a small fraction of synaptosomes (∼3%) from control brains, reaching ∼20% by Braak stage 6 with more larger aggregates. We then investigated the multi-phosphorylation of synaptic tau aggregates for AT8 and T181 and quantified the co-localisation of phosphatidylserine and CD47, synaptic “eat me” and “don’t eat me” signals respectively, along with synaptogyrin-3, which contributes to tau mediated synaptic dysfunction. T181, phosphatidylserine, and synaptogyrin-3 co-localisation with AT8-positive tau were increased during stage 3 and CD47 was decreased, indicating early synaptic pathology is associated with the formation of small tau aggregates, contributing to microglia-driven synaptic loss.

## 1. Introduction

Alzheimer’s disease (AD) is the globally leading cause of dementia, with the primary symptoms of cognitive impairment and memory loss^1^. While the extracellular beta-amyloid (Aβ) plaques and neurofibrillary tangles made of hyperphosphorylated tau are the neuropathological hallmarks of AD, synaptic dysfunction and loss correlate with cognitive decline more strongly than plaque/tangle formation and neuronal loss^2,3^. Noteworthily, synaptic dysfunction has been reported to precede the Aβ plaques and neurofibrillary tangles in AD brain and animal models^4,5^, raising the question of the temporal order of events in AD. Recently it has been proposed that the small aggregates of Aβ and tau are highly toxic. These small aggregates are described as “diffusible” since they remain in the supernatant after a high-speed centrifugation, separating them from the fibrils making up the plaques and tangles^6,7^. As they are capable of interacting with various receptors^8^, organelles^9,10^, and nucleic acids^11,12^, they have been suggest to be the primary species promoting AD pathology.

While the presence of small-diffusible (sometimes termed oligomeric) tau aggregates has been documented in the synapses of the AD brain along with fibrillar tau aggregates^13–15^, the patho-physiological outcome(s) of the accumulation of these species at synapses is not fully understood. One of the prominent theories related to the synaptic accumulation of tau is that this triggers microglia activation, which then prune the synapses and acquire a disease-associated highly inflammatory phenotype^16,17^. These disease-associated microglia then further promote AD pathology through inflammatory mechanisms^18,19^. Another proposed disease mechanism involves the direct synaptic transmission of tau aggregates which has been documented in mouse models^20–22^ and suggested from functional-connectivity studies in humans^23,24^. Since aggregated tau is known to have seeding activity^25^, it has been proposed that the synaptic transmission of tau aggregates may cause seeding-driven aggregation in functionally connected neurons, in a “prion-like” manner^26–28^. While there is evidence for both the inflammation-driven (indirect) and seeding-driven (direct) spread of tau pathology in AD, the relative contribution of these mechanisms and their relationship with oligomeric and fibrillar tau is an active area of research. Although some studies did suggest the synaptic transmission^29^ and seeding competence of oligomeric tau^30^, seeding is primarily studied using fibrillar aggregates^31,32^, and it has been shown that fibrillar aggregates of tau around 170 nm in length are the most seeding-competent species^33^. Meanwhile, the smaller and rounder, “oligomeric” tau aggregates in synapses are shown to induce local inflammation through microglia activation^30^ which was also shown to be induced by fibrillar tau presence in the neurites^33^. As such, even though neither mechanism exclude the probability of the other and ongoing studies provide evidence for both to likely occur in the same brain, these pathological outcomes of synaptic tau aggregate accumulation could be associated with different types of tau aggregates. This link between tau aggregate morphology and pathological outcome makes it crucial to further characterise the actual tau aggregates present in the synapses at different disease stages, so appropriate therapeutics can be developed targeting the “right” aggregates. Indeed, recent studies on Aβ aggregates have shown the importance of targeting the morphologically correct species for therapeutic success^34,35^.

While the evidence for the presence of different types of tau aggregates in the synaptic compartments and their role in synaptic and cellular dysfunction is growing, the small size of these aggregates (which is mostly below the diffraction-limit of light) along with their low abundance, and high heterogeneity of their morphology and post-translational modifications makes them challenging to detect and characterise at single-synapse level. Our group has developed advanced single-molecule detection tools using super-resolution microscopy^36–38^ and digital-ELISA^39^ based methods to quantify and characterise the small aggregates formed in neurodegenerative diseases and we have recently extended these techniques to synaptosomes harvested from post-mortem human brains and mouse models. This technique, called SynPull, combines single-molecule pulldown and single-molecule microscopy to quantify and characterise small aggregates inside individual synaptosomes^40^. Using SynPull, we have investigated the Aβ and AT8-positive^41^ tau aggregates in late stage-AD^40^ and mouse models, as well as alpha-synuclein aggregates in Parkinson’s disease brain samples and mouse models^40^. These analyses demonstrated that small tau aggregates are the dominant species in the pre-synaptic compartment in AD, indicating their role in synaptic pathology.

In the current study, we utilised SynPull to investigate synaptic tau aggregation at different stages of AD (Braak stage 0, 3, and 6) in human post-mortem pre-frontal cortex samples, characterising the synaptic and extra-synaptic AT8-positive tau aggregates throughout the disease progression and particularly at early stages. Braak stages 3 and 6 were chosen to study the synaptic alterations prior to (Stage 3) and during the presence of (Stage 6) tau tangle pathology in the pre-frontal cortex^42^, in order to investigate the temporal order of small tau aggregate accumulation in the synapse and insoluble tangle formation. Using direct stochastic optical reconstruction microscopy (*d*STORM) imaging, we were able to quantify and characterise individual aggregates inside synaptosomes, in terms of their size and shape, providing accurate data on their morphology at single-aggregate and single-synapse level. Then, we investigated multi-phosphorylation of tau at the signal synapse-level, by co-localising AT8 and T181 phosphorylation. Lastly, we quantified the co-localisation of (patho)physiological markers phosphatidylserine^43^, CD47^44^, and synaptogyrin-3^45^, with AT8-positive tau, in individual synaptosomes, determining the probability of finding these markers alongside small tau aggregates in individual synaptosomes. Although our previous results did not show significant Aβ aggregate accumulation in AD synapses, diffusible Aβ aggregates can still alter synaptic physiology through proteomic changes and by inducing the translocation of tau to the synapse^46,47^. As such, we have also quantified the diffusible Aβ levels in the same brain samples, to investigate their possible association with synaptic tau pathology (**Supplemental Figure 1A&B**). Collectively, our results suggest that synaptic tau aggregation starts early in AD by Braak stage 3 -preceding neurofibrillary tangle formation in the pre-frontal cortex but during the presence of small-diffusible Aβ aggregates, and progresses with disease stages, with the majority of the aggregates remaining non-fibril-like. Moreover, the presence of tau aggregates in individual synapses alters the presence of markers associated with synaptic dysfunction and loss -linking small tau aggregates to microglia-driven synaptic pathology in AD.

## 2. Results

### Synaptic tau aggregation in AD

To characterise the small tau aggregates positive for AT8 phosphorylation (referred as AT8+ aggregates), we used SynPull (**Figure 1A**) on the AD pre-frontal cortex samples. While the presence of AT8+ tau in the synapse was not AD specific and also observed in control (Braak stage 0) cases, both the fraction of synaptosomes with aggregates (**Figure 1B**) and the number of aggregates inside these synaptosomes (**Figure 1C**) were significantly less compared to disease samples. The fraction of synaptosomes containing one or more AT8+ tau aggregate showed a Braak stage-dependent increase, with an initial rise from 2% to 13% at stage 3 and a further increase in stage 6, reaching 20% (**Figure 1B)**. The number of aggregates per synapse showed a similar trend with an early increase at stage 3 and differed significantly between stages 0 and 6 (**Figure 1C**). After quantifying them, we characterised the small tau aggregates in terms of their size and shape using *d*STORM imaging. While the length of the synaptic AT8+ tau aggregates also increased with disease stage, with an average length of 117 nm at stage 0, 154 nm at stage 3 and 182 nm at stage 6 (**Figure 1D**), they mostly remained non-elongated (circular) with average eccentricity values remaining below 0.8 (**Figure 1G**). When we focused on the longer aggregates above 150 nm in length, the average aggregate eccentricity decreased with Braak stage (**Figure 1H**). Large and elongated aggregates with a length over 150 nm and eccentricity over 0.9 remained rare yet significantly higher in AD cases, with a 6% occurrence rate in the control cases, 16.5% at stage 3, and 15% at stage 6 (**Figure 1I**). Collectively these results show that synaptic aggregation of AT8+ tau begins early in AD and affects more synapses as the disease progresses, accompanied by morphological changes in the aggregates as they grow in size but mostly remain non-elongated.

**Figure 1.**
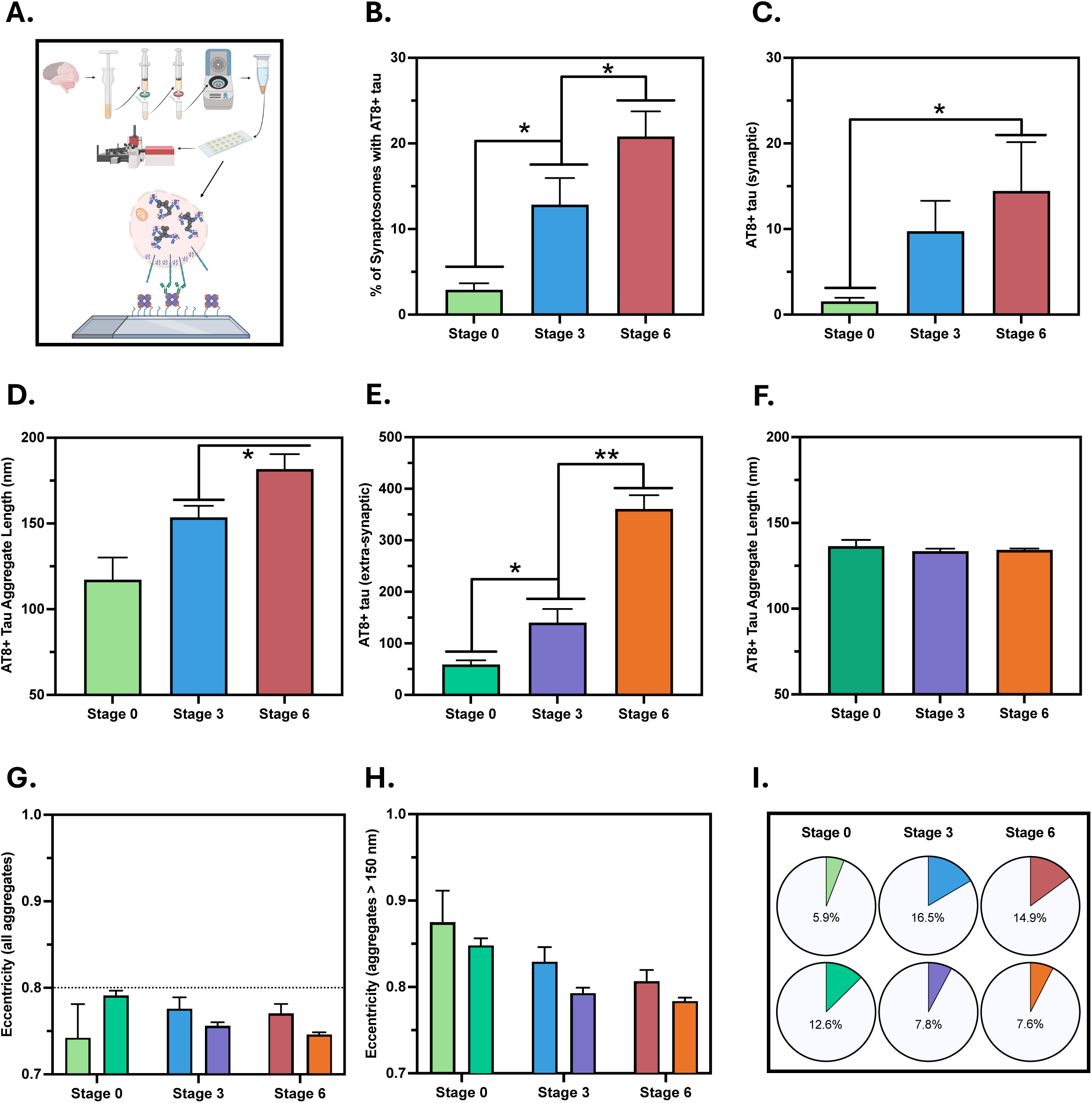
Synaptic AT8-positive tau aggregate quantity and morphology determined using SynPull. (**A**) Cartoon representation of SynPull methodology, showing the homogenisation, filtration, centrifugation, and imaging steps. (**B**) Percentage of synaptosomes carrying one or more aggregate, (**C**) number and (**D**) length of the aggregates within the synaptosomes, (**E**) number and (**F**) length of extra-synaptic aggregates at each Braak stage. (**G**) Eccentricity of the synaptic aggregates, indicating their shape with 0 being a perfect circle and 1 being a flat line. (**H**) Eccentricity of the synaptic aggregate above the length of 150 nm. (**I**) Fraction of synaptic (Syn) and extra-synaptic (Extra-syn) aggregates at each Braak stage with a length over 150 nm and eccentricity above 0.9. These aggregate are characterised as “fibril-like”. Date in B-H are represented as Mean±SEM and differences are calculated using 95% confidence intervals (CI) with a CI not including 0 is considered significant, indicated with an asterisk.

On the other hand, quantity of the extra-synaptic aggregates showed a much greater increase from Braak stage 3 to 6, following a smaller change between stages 0 and 3 (**Figure 1E**) However, the length of the extra-synaptic AT8+ tau aggregates did not show an AD stage-related difference and remained constant with a mean of approximately 135 nm (**Figure 1F**). Similar to the synaptic aggregates, the extra-synaptic AT8+ tau aggregates were also mostly round, with even smaller eccentricity compared to synaptic aggregates (**Figures 1G-H**). Interestingly, while the fraction of long and elongated aggregates were higher for the extra-synaptic aggregates in the control cases (6% synaptic vs 13% extra-synaptic), this trend reversed in AD with 7-8% of the extra-synaptic tau aggregates longer than 150 nm in length with an eccentricity greater than 0.9 (**Figure 1I**). Taken together, these results show that the number of small tau aggregates outside the synapse increases at later stages of AD, but no significant changes in aggregate length occur. This contrasts the early rise in synaptic tau aggregates, which continue to grow is size, suggesting that the synapse may be a more susceptible region of aggregation. This was also shown by analysis of the aggregate length distribution. Both the synaptic and extra-synaptic aggregate length distributions were consistent with a geometric decay, which results from the balance of elongation and removal processes^48^. However, the synaptic aggregate length distribution decays more slowly with increasing length, indicating that the balance of elongation and removal processes inside the synapse is shifted more in favour of aggregation. We quantify the ratio of removal rate to elongation rate and find this to be reduced by 45±6% for aggregates from synapses compared to the extra-synaptic ones (**Figure 2**). While majority of the AT8+ tau aggregates in AD are round, there is a greater fraction of elongated aggregates in the synapse than in the extra-synaptic fraction, showing distinct tau aggregate patterns between different parts of the neurons. The low concentration of elongated aggregates and abundance of small-aggregates, which has been associated with local inflammatory responses^30^ suggest that local inflammation may play a key role. To further study this, we looked at synaptic markers of microglia activation and synaptic function.

**Figure 2.**
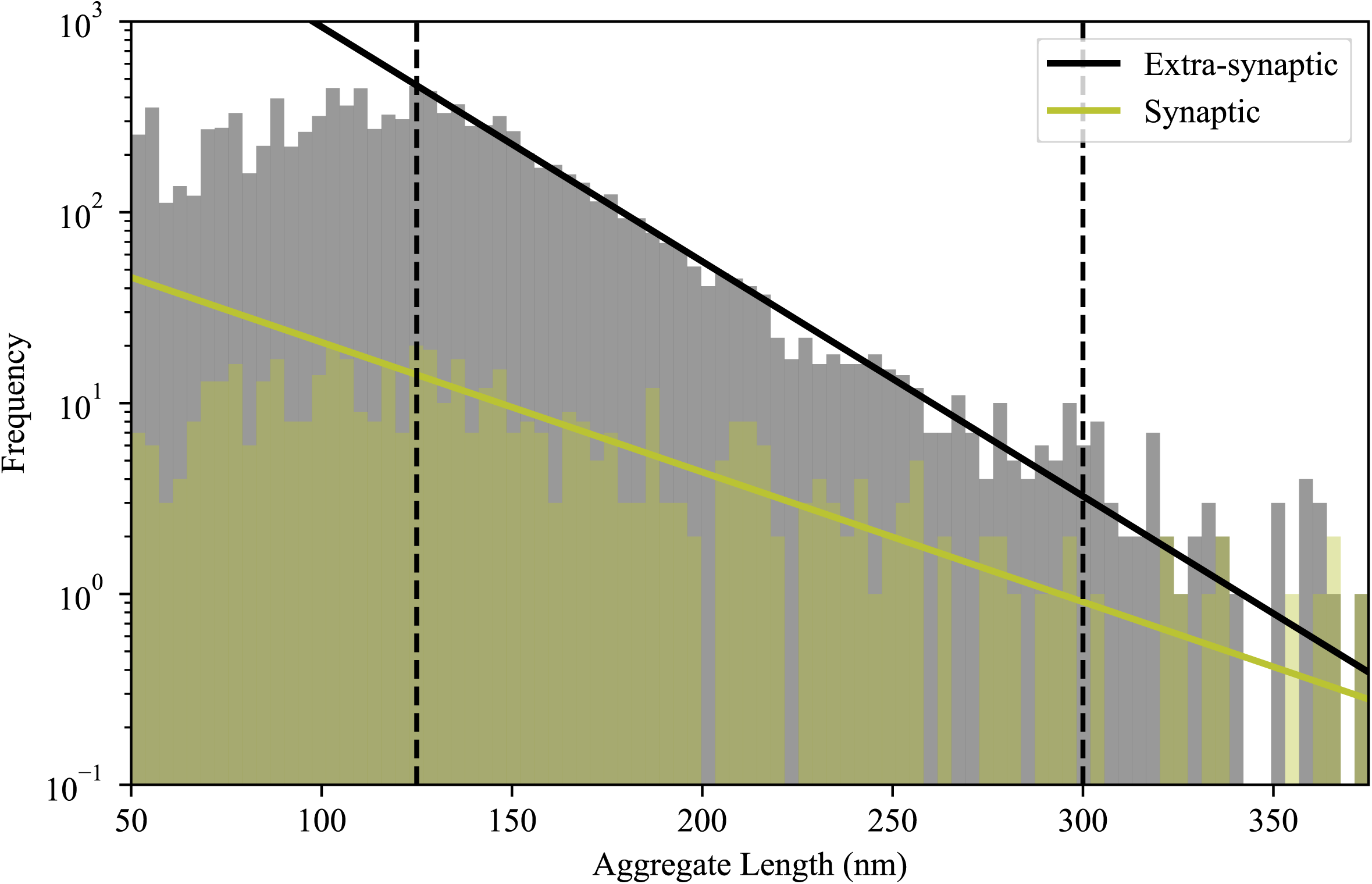
Histogram of the synaptic and extra-synaptic aggregate length distributions (in nanometres (nm)) across all Braak stages. The solid lines show the maximum likelihood fit of equation (1) to the aggregate populations in each region. The two dashed lines show the upper and lower bounds of the fitted region.

### Synaptic tau becomes multi-phosphorylated and co-localises with functional markers

After showing the presence of AT8+ tau aggregates in the synapse at early stages of AD, that are mostly round and not elongated, we investigated the physiological and pathological outcomes that correlate with this accumulation. Studying individual synaptosomes and co-localising the AT8+ tau signal with T181-positive tau at the aggregate level, as well as determining the presence of exposed phosphatidylserine (“eat me” signal), CD47 (“don’t eat me” signal), and synaptogyrin-3 in synaptosomes containing AT8+ tau, we were able to explore how synaptic tau aggregation alters synaptic function and pruning, which are potentially contributing to cognitive deficits in AD. In order to do this, we determined 4 parameters: (1) the total amount of signal in the FoV, (2) the percentage of the synaptosomes carrying the signal ([Synaptosomes with Signal) / All Synaptosomes]*100) -which estimates the marginal probability of observing the signal, (3) the percentage of the synaptosomes carrying the signal as well as AT8 out of the number of synaptosomes carrying AT8 ([Synaptosomes with Signal + AT8+ tau) / Synaptosomes with AT8+ tau]*100) -which estimates the conditional probability of observing the signal given the presence of AT8+ tau, and (4) the percentage of the synaptosomes carrying the signal as well as AT8 out of the total number of synaptosomes ([Synaptosomes with Signal + AT8+ tau) / All Synaptosomes]*100) -which estimates the joint probability of observing both AT8+ tau and the signal. Based on these data, we used Bayesian inference^49^ to estimate the probabilities of co-occurrence of each type of signal with AT8+ tau, and compared this to the probability of occurrence of the signal regardless of the presence of AT8+ tau. This allowed us to determine if the presence of AT8+ tau lead to a changed probability of observing the signal, and how this correlation varies over disease stages.

While the total T181-positive tau signal in the sample (**Figure 3A**) and the fraction of synaptosomes with T181-positive tau (**Figure 3B**) were both significantly higher in the Braak stage 3 samples compared to the controls, neither value was further elevated at stage 6. Similarly, multi-phosphorylation at AT8 and T181 as a fraction of synaptosomes with AT8+ showed a significant difference only between stages 0 and 3 (**Figure 3C**), however the synaptosomes carrying multi-phosphorylated aggregates as a fraction of all synaptosomes (with or without AT8+ tau), showed a significant increase at stage 3 from stage 0 and a further increase at stage 6 (**Figure 3D**). Together, these results show that tau phosphorylated at T181 increases early in AD and multi-phosphorylated tau is increased in disease and exists in more synapses as the disease progress.

**Figure 3.**
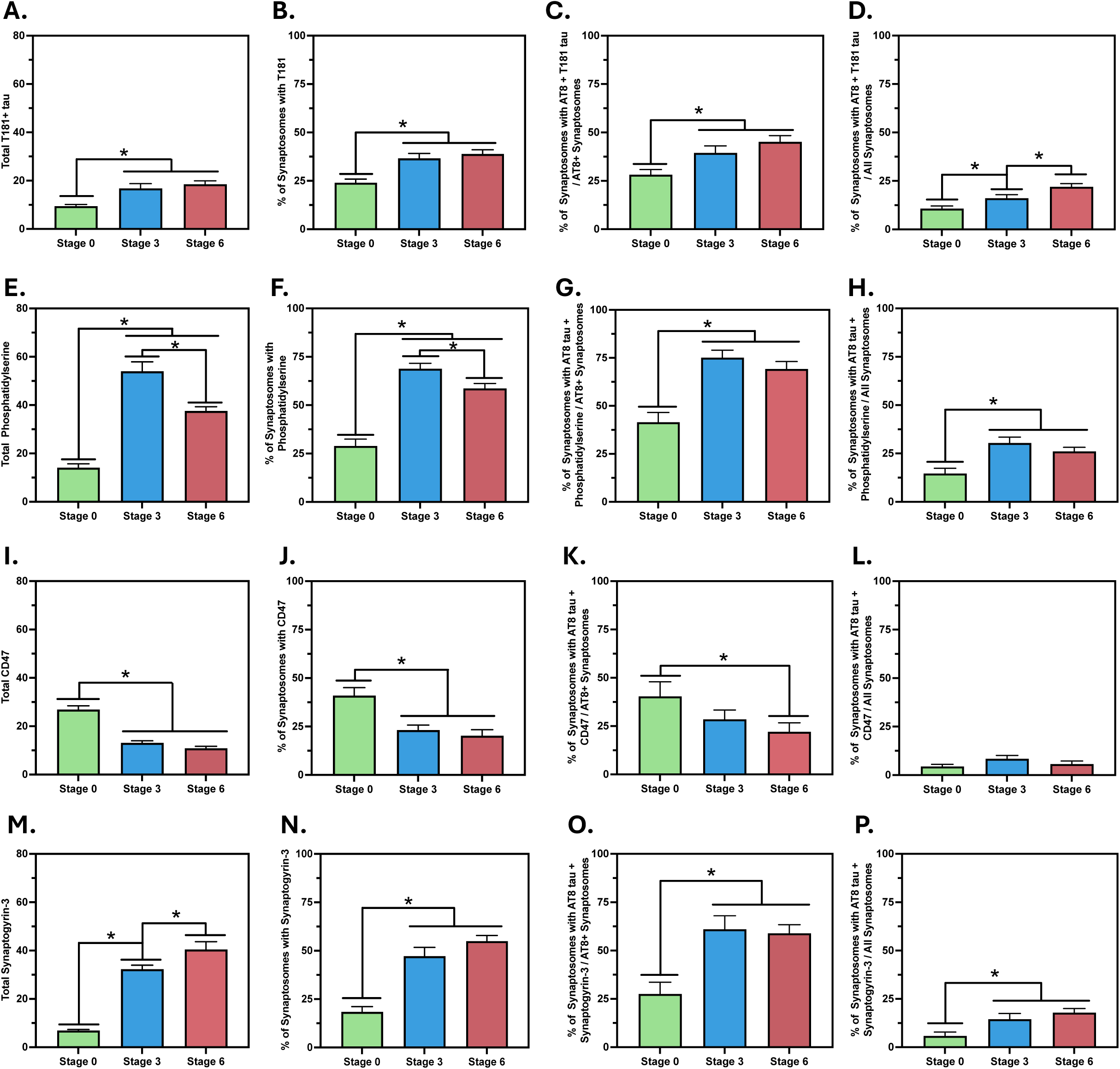
Pathology-associated markers measured inside the synaptosomes and co-localised with AT8-positive tau aggregates. (**A/E/I/M**) Total number of T181-positve tau (**A**), phosphatidylserine (**E**), CD47 (**I**), and synaptogyrin-3 (**M**) per field of view (FoV). (**B/F/J/N**) Synaptosomes carrying a T181-positve tau (**B**), phosphodiesterase (**F**), CD47 (**J**), and synaptogyrin-3 (**N**). (**C, G, K, O**) Percentage of synaptosomes carrying a T181-positve tau (**C**), phosphatidylserine (**G**), CD47 (**K**), and synaptogyrin-3 (**O**) and AT8-positive tau (co-localised in the same synaptosome), as a percentage of synaptosomes with AT8-positive tau. (**D, H, L, P**) Percentage of synaptosomes carrying a T181-positve tau (**D**), phosphodiesterase (**H**), CD47 (**L**), and synaptogyrin-3 (**P**) and AT8-positive tau (co-localised in the same synaptosome), as a percentage of all synaptosomes. Date in are represented as Mean±SEM and differences are calculated using 95% confidence intervals (CI) with a CI not including 0 is considered significant, indicated with an asterisk.

Then, we investigated the “eat me” signal phosphatidylserine, which contributes to microglial synaptic pruning when exposed on the extracellular side of the cell membrane^43^. By using an antibody that specifically recognises exposed phosphatidylserine, we showed that while the total phosphatidylserine levels were increased at stage 3, they showed a reduction at stage 6 compared to stage 3 (**Figure 3E**). Similarly, the fraction of synaptosomes with exposed phosphatidylserine also showed an early increase at stage 3 followed by a reduction at stage 6 (**Figure 3F**). Together, these results indicate an early increase in microglia-mediated synaptic pruning signalling in AD, which decreases in the later disease stages. When we co-localised the “eat me” signal with AT8+ tau in the synaptosome, the percentage of co-localisation within the synaptosomes with AT8+ tau was significantly higher at stage 3, compared to stage 0, but did not further change at stage 6 (**Figure 3G**). Similarly, the synaptosomes with the co-localisation of phosphatidylserine and AT8+ tau as a percentage of all synaptosomes was increased at stage 3 but did not show any further changes at stage 6 (**Figure 3H**). Bayesian inference demonstrated a clear correlation between the presence of AT8+ tau and the presence of phosphatidylserine, with a significantly higher probability of observing the signal when AT8+ tau was present, across all stages of the disease. As demonstrated by the very small overlap of the posterior probabilities in **Figure 4A**, the difference was significant across all stages. Specifically, the probability that AT8+ tau has an effect on the presence of an eat-me signal was calculated as 99.9% in Braak stage 0, 99.1% in stage 3 and 85.1% in stage 6. Collectively, these results suggest an increase in phosphatidylserine exposure early in AD, possibly regulated by synaptic tau aggregation through interaction with flippases regulating the membrane localisation of phosphatidylserine^50^ or by directly interacting with phosphatidylserine^51^. However, this is not a progressive condition as the levels are not further increased in later AD, suggesting additional mechanisms may be contributing to synaptic loss at later disease stages.

**Figure 4.**
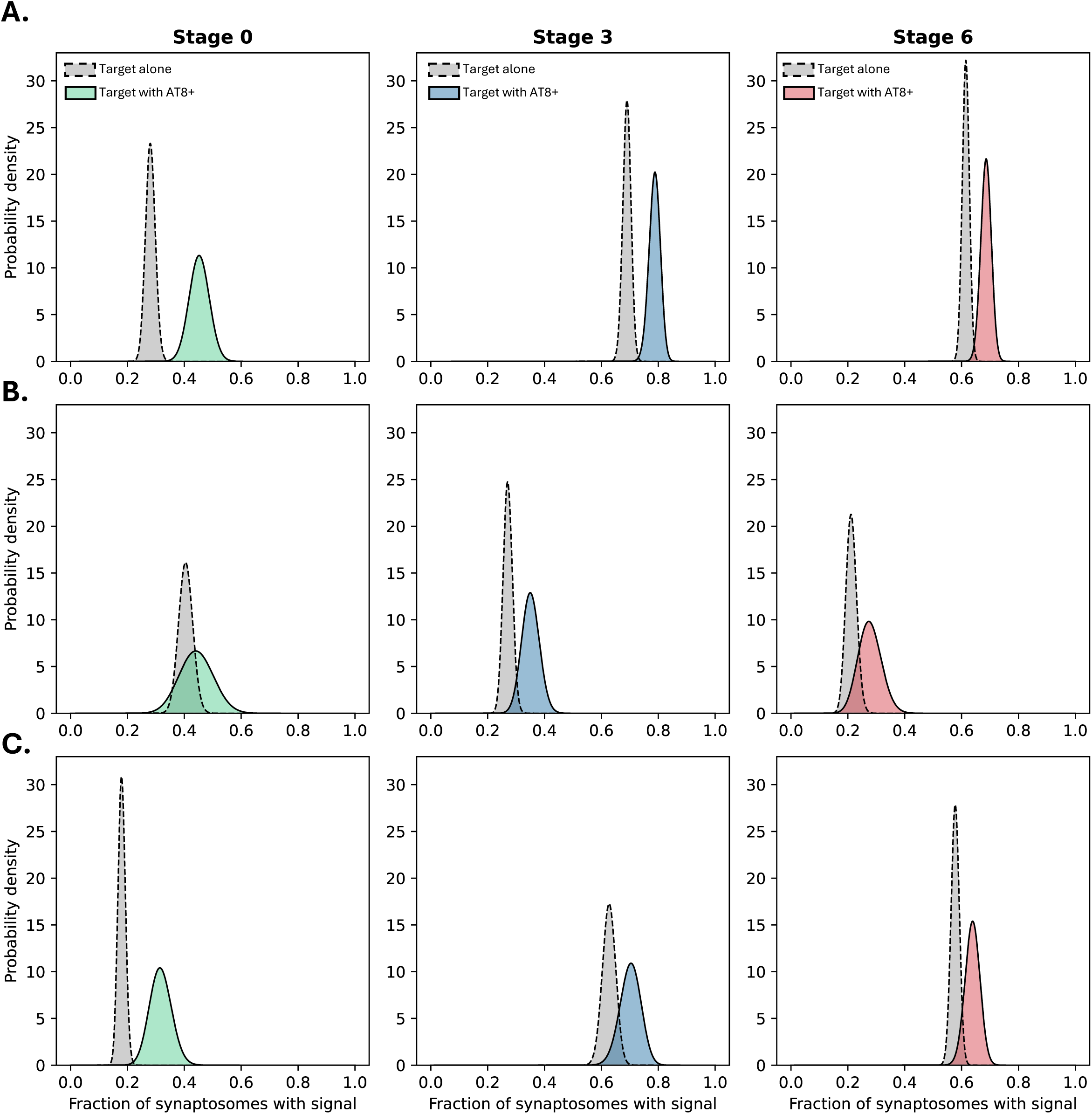
Probability to encounter synaptic markers and its correlation with AT8 state: Bayesian Inference was performed to estimate the probability of encountering each of the synaptic markers, (**A**) phosphatidylserine, (**B**) CD47, and (**C**) synaptogyrin-3. The posteriors are shown for encountering the signal regardless of AT8 state, i.e. the marginal probability (grey curves), and for encountering the signal if the synaptosome is AT8 positive, i.e. the conditional probability (coloured curves). A clear separation of the posteriors implies a significantly increased probability of encountering the signal in AT8 positive synaptosomes, as can be seen across stages for phosphatidylserine.

Following the “eat me” signal, we focused on the “don’t eat me” signal CD47 in terms of its relationship with AT8+ tau at the single-synapse level. While the total CD47 levels in the samples were significantly reduced at Braak Stage 3 compared to Stage 0, there was no difference between Stage 3 and 6 (**Figure 3I**). In parallel to total CD47, the percentage of synaptosomes carrying the “don’t eat me” signal was also reduced significantly by Stage 3, without any further reduction at Stage 6 (**Figure 3J**). When the co-localisation of AT8+ tau and CD47 within synaptosomes with AT8+ tau was compared between the Braak stages, no difference was seen between Stage 0 and Stage 3 or Stage 3 and 6, yet the fraction of synaptosomes with CD47 and AT8+ tau co-localisation was significantly lower in stage 6 AD brains compared to the stage 0 controls brains (**Figure 3K**). Meanwhile, the fraction of synaptosomes with the co-localisation of CD47 and AT8+ tau within all synaptosomes did not show any significant difference between disease stages (**Figure 3L**). Collectively, these results show that while the expression of CD47 is reduced in AD in a progressive manner, the co-localisation of this “don’t eat me” signal and AT8+ tau is a rare event in the synaptosomes regardless of AD status. This was also evident from the strong overlap of posterior probability distributions in **Figure 4B**, as the presence of AT8+ tau correlated only weakly with CD47, with the probability that AT8+ tau impacting the presence of CD47 in the same synaptosome calculated as 10.9% in Braak stage 0, 44.7% in Stage 3 and 16.7% in Stage 6, suggesting that pathological tau induced loss of CD47 at the single synapse level may not be a primary mechanism of synaptic loss in AD.

Lastly, we quantified synaptogyrin-3 -which has been reported to mediate the binding of tau aggregates to neurotransmitter vesicles and cause pre-synaptic dysfunction and investigated its co-localisation with AT8+ tau in the synaptosomes^45^. Total synaptogyrin-3 levels were significantly higher in Braak stage 3 compared to Stage 0 and in Stage 6 compared to stage 3 (**Figure 3M**). Meanwhile the fraction of synaptosomes containing synaptogyrin-3 were elevated at stage 3 but did not further change at stage 6 (**Figure 3N**). Similarly, when compared between the Braak stages, the percentage of synaptosomes with a co-localisation of synaptogyrin-3 and AT8+ tau, within the synaptosomes with AT8+ tau also showed an increase in Stage 3 yet did not differ between Stage 3 and 6 (**Figure 3O**). When the percentage of synaptosomes with co-localisation of synaptogyrin-3 and AT8+ tau was calculated within all synaptosomes, once again increased levels were observed at Stage 3 without further changes in Stage 6 (**Figure 3P**). Collectively, these results show an early increase in synaptogyrin-3 in AD, contributing to synaptic dysfunction. According to Bayesian inference, the probability of observing synaptogyrin-3 was increased in AT8+ tau containing synaptosomes. However, a strong correlation was observed only at Stage 0, with weaker correlations at the later stages (**Figure 4C**). Specifically, the probability that AT8+ tau has no effect on the appearance of Synapotgyrin-3 was 97.0% in Stage 0, 27.6% in Stage 3, and 29.6% in Stage 6.

## 3. Discussion

The role of synaptic tau aggregation on the morphological and (patho)physiological alterations in synapses is an active area of research^13,15,22^. While physiological presence and activity of tau has been reported in the synapse^52^, it has also been shown that oligomeric Aβ can initiate the pathological translocation and aggregation of tau to the synapse leading to deficits^47^. Synaptic tau aggregates can interact with pre-synaptic vesicles to cause neuronal communication deficits^53^ and alter the synaptic proteome^54^, leading to synaptic dysfunction and loss, chiefly regulated by microglia^17^. Intriguingly, it was recently shown that in AD patients who showed behavioural symptoms of dementia and brains showing Braak stage 3 and 5 pathology, small tau aggregates are present in the synapses at a greater extent compared to patients with the same neuropathological features (same Braak staging) who were resilient to dementia symptoms^14^, suggesting that the synaptic localisation of small tau aggregates is a major driver of cognitive decline in AD. In order to further investigate this disease mechanism, we studied synaptic tau aggregates at Braak stage 0, 3, and 6, post-mortem human pre-frontal cortex samples using SynPull, which is a method we have developed^40^ that allows single-molecule investigation in individual synaptosomes. We have also quantified the co-localisation of synaptic proteins associated with pathology and protection, to understand the patho-physiological outcomes of the synaptic small tau aggregate accumulation.

We used *d*STORM super-resolution imaging to quantify and characterise the synaptic and extra-synaptic AT8+ tau aggregates in the same pre-frontal cortex samples. While the synaptic aggregates showed an early increase in quantity by stage 3, their numbers did not increase significantly by stage 6, yet the fraction of synapses containing aggregates and the size of the aggregates inside the synaptosomes continued to grow by stage 6, showing continuous aggregation throughout the progression of AD. On the other hand, there was no size difference for the extra-synaptic tau aggregates between the Braak stages, but their quantity increased dramatically after stage 3, following a relatively miniscule increase at stage 3. This suggests that synaptic aggregation of AT8+ tau is not only an early event in AD, but it also precedes the pathological aggregation seen in the rest of the neuron. While this may be due to the active translocation of tau to the synapse by the Aβ aggregates^47^, oligomeric tau aggregates in the synaptosomes were also found to be hyperphosphorylated and associated with ubiquitinated substrates and elevated proteasome components^13^, suggesting that the post-translational modifications of tau and a reduction in its aggregate removal rate would also contribute to this early synaptic aggregate accumulation. Strikingly, only a small portion of the AT8+ tau aggregates in the synaptic and extra-synaptic fractions were longer than 150 nm in length and showed high eccentricity, suggesting the lack of an elongated morphology (**Figure 1I**), but the concentration of these species were higher in the synaptic fraction of AD cases (around 15%). By comparing the length distribution of synaptic and extra-synaptic aggregates, we were able to quantify the relative rate of aggregate removal compared to aggregate growth. We found that the longer aggregates encountered in the synapse indicate that the relative rate of removal there is approximately half the rate for extra-synaptic aggregates, providing an explanation for earlier aggregation in the synapse and the early synaptic deficits in AD. Although it should be noted that the source of the extra-synaptic aggregates are not only the rest of the neuron, as they may also be harvested from glial cells or the extra-cellular space. Spreading of tau pathology through functionally connected brain regions in a “prion-like” manner has been suggested^20,22,23^ and fibrillar tau aggregates around 170 nm in length have been demonstrated to be the most seeding-competent species^28,31–33^. On the other hand, the smaller oligomeric tau aggregates are shown to promote local inflammation driven by a microglia response^30^. Our results do not rule out synaptic spreading of tau pathology through functional connectivity and seeding as a relevant mechanism as we do observe a small amount of aggregates associated with this disease mechanism, specifically in the AD cases. However, the fact that the majority of the aggregates are small and non-elongated supports the idea that local microglia response to synaptic tau aggregates and the resulting inflammation play a key role in driving AD pathology. This interpretation is further supported by the association of microglia-response related signals with tau aggregates inside individual synapses, as further discussed below.

In order to check the phosphorylation state of the synaptic tau aggregates, we quantified the co-localisation of AT8- and T181-positive tau signal in individual synaptosomes. The fraction of synaptosomes with T181-positive tau as well as synaptosomes with co-localisation, as a fraction of all synaptosomes with AT8+ tau were increased by stage 3, yet did not change by stage 6, showing that multi-phosphorylation of tau in the synapse is indeed an early event in AD. Nevertheless, the AT8 and T181 co-localisation showed a further increase in stage 6, showing that the newly formed aggregates inside the synapses at later stages of AD are more likely to be multi-phosphorylated. These results agree with the previous findings of hyperphosphorylated tau aggregates being present in the synaptosomes in AD^13^.

After showing the presence of tau aggregates in synaptosomes as an early event in AD pathogenesis, we investigated the patho-physiological outcomes of this accumulation. A number of proteins have been associated with synaptic dysfunction and synaptic pruning, including phosphatidylserine, CD47, and synaptogyrin-3. Phosphatidylserine is also known as the “eat me” signal, as it mediates microglial synaptic pruning, regulated by TREM2^43^. Our previous work has shown that neurons with tau filaments expose phosphatidylserine before getting phagocytosed by microglia^55^ and spines that become overactive through oligomeric Aβ stimulation get pruned by microglia upon externalising phosphatidylserine^56^. On the other hand, CD47 works as a “don’t eat me” signal, protecting synapses from microglia-mediated pruning during development^44^. Meanwhile, pre-synaptic co-localisation of tau with synaptogyrin-3 contributes to synaptic plasticity deficits along with working memory dysfunction and genetic silencing of synaptogyrin-3 ameliorates these symptoms in mouse models of tau pathology, without altering tau induced neuroinflammation^57^. We quantified these markers in individual synapses as well as their co-localisation with AT8+ tau, which allowed us to determine how the probability of the presence of these signals is altered by the presence of AT8+ tau.

It has previously been shown that tau oligomer-containing synapses in AD are eliminated by microglia and astrocytes^14^. In accordance with these findings, phosphatidylserine signal showed a significant increase with early disease stages and Bayesian inference showed a higher chance for the presence of this signal in AT8+ tau containing synaptosomes, even in non-AD brain samples and late-stage AD, where the fraction of synaptosomes containing phosphatidylserine was reduced. Although a causal mechanism needs to be further demonstrated in future studies, this finding indicates a role of synaptic tau aggregate accumulation and “eat me” signals in tau-driven early synaptic loss in AD.

Through a similar mechanism, phosphatidylserine also works as an “eat me” signal during developmental synaptic pruning^43^, while this signal is counteracted by CD47, which protects the synapses^44^. Interestingly, increased CD47 levels were observed in the AD hippocampus, co-localising with synaptic tau aggregates^58^. While our analysis in the pre-frontal cortex samples showed a Braak-stage dependent decrease in overall CD47 levels, there was a weak trend of increase in the presence of CD47 in synaptosomes that are AT8+ tau at Braak stage 3. These findings suggest that while synaptic protective signals are reduced in AD, localising a “don’t eat me” signal in the affected synapses may be an attempt by the neurons to reduce synaptic loss during early stages, when tau-mediated synaptic loss becomes an active disease mechanism.

We investigated synaptic dysfunction caused by tau aggregation, by characterising the co-localisation of synaptogyrin-3 with AT8+ tau in individual synapses. Similar to the “eat me” and “don’t eat me” signals, synaptogyrin-3 was also increased in AD, yet Bayesian inference did not show a meaningful effect of AT8 on increasing the presence of synaptogyrin-3 at the disease samples. Synaptogyrin-3 mediates the binding of tau to pre-synaptic vesicles^45^, which causes deficits in vesicle mobility and neurotransmitter release^53^. This suggests indicates a pathological role of synaptogyrin-3 in relation to the synaptic presence of tau, by mediating its interaction synaptic vesicles. As such, the presence and increase of synaptogyrin-3 in the synapse may be an independent event from synaptic tau aggregation and even precede it. In agreement with this, increased synaptic synaptogyrin-3 levels, compared to stage 0, were observed at stage 3 and 6. Since this increase was not linked to tau aggregation, there may be another, upstream mechanism. Activation of nuclear receptor 4A2 (NR4A2, also known as NURR1) was shown to increase synaptogyrin-3 expression^59^ and Moon et al.,^60^ have shown a significant positive correlation between NR4A2 expression and Aβ aggregation in the 5xFAD mouse model of Aβ accumulation. This association between Aβ, NR4A2, synaptogyrin-3, and tau aggregates propose an AD mechanism, with which Aβ aggregation leads to higher synaptogyrin-3 levels, which contribute to tau mediated synaptic dysfunction. Indeed, we have shown (**Supplemental Figure 1B**) an early increase in stage 3 -preceding an increase in total soluble Aβ levels, further supporting the presence of this AD mechanism.

### Conclusions, limitations, and future directions

In this study we investigated the synaptic accumulation of small tau aggregates and their possible role in synaptic dysfunction through various markers in post-mortem AD pre-frontal cortex samples at different Braak stages. While the synaptic accumulation of AT8+ tau aggregates was already apparent at early stages of AD, morphology of these aggregates continued to evolve by Braak stage 6, showing that synaptic tau aggregation remains a disease process throughout the pathogenesis of AD. Importantly, the presence of small-diffusible Aβ aggregates correlated with synaptic tau pathology, suggesting possible links, such as synaptogyrin-3 mediated tau binding to synaptic vesicles. On the other hand, synaptic pruning seems to be directly mediated by synaptic tau aggregation, by an increase of phosphatidylserine, which functions as an eat me signal for microglia, as the presence of this signal was predicted by Bayesian inference to be directly linked to synaptic AT8+ tau aggregates. Meanwhile, there seems to be an attempt of reducing this synaptic loss, with an increase of CD47 signalling, which works as a “don’t eat me signal” during neurodevelopment. It is noteworthy that the “eat me” and “don’t eat me” signals are important for brain development at early stages of life are also involved in AD, much later on, showing that AD hijacks the homeostatic physiology of the nervous system during its pathogenesis, rather than using novel mechanisms. Importantly, all of the pathological changes we studied here were most prominent at Braak stage 3, before an exponential increase in extra-synaptic tau aggregate levels and tau tangle pathology in the pre-frontal cortex, supporting that early synaptic deficits precede neuronal dysfunction and loss in AD, and are mediated by small-diffusible aggregates rather than much larger plaques and tangles. While some aggregates longer than 150 nm in length with fibril-like morphologies were detected, the very low concentration of these aggregates relative to the shorter and rounder species indicate that these smaller aggregates, and thus the related microglial response, play a more prominent role in synaptic dysfunction during the early stages of AD.

It must be noted that the synaptosome extraction protocol we use in this study predominantly harvests the pre-synaptic compartment with parts of the post-synapse attached in some cases^40,61^ and thus the post-synaptic tau aggregates cannot be characterised. Since the post-synaptic accumulation of tau has been shown in AD^22^ and the synaptic spread of tau in functionally connected neurons has been suggested as a disease mechanism^23^, the inability to study post-synaptic tau aggregates and their co-localisation with markers associated with post-synaptic dysfunction is a limitation of this work. Moreover, tau leads to synaptic loss and thus there is a survival bias, through which the (patho)physiology of the synapses that remained intact during the time of death of the AD patient may differ from ones that were lost throughout the disease process. Lastly, since we only studied post-mortem human brain samples, while our results are highly valid, they remain correlational. Future work on animal and cellular models with direct manipulation of tau as well as other markers that we have studied here will be beneficial for identifying the direct and temporarily causal links between synaptic tau aggregation, pathology, and cognitive decline. Nevertheless, the results we present here provide strong support for the early synaptic aggregation of multi-phosphorylated tau aggregates that are small in size (below the diffraction-limit of light) and involved in microglia mediated synaptic pruning by altering the levels of markers also involved in early neurodevelopment.

## 4. Methods

### Human brain samples and synaptosome preparation

15 post-mortem brain samples were used in the experiments, with Braak stage 0, 3, and 6 (5 samples each) pathology (**Table 1**). Samples were collected from the orbito-frontal cortex (Brodmann’s area 10-11) and snap frozen and stored frozen at –80□ until synaptosome preparation.

**Table 1.** Demographic data for human frontal cortex tissue samples.

| **Case** | **Age** | **Sex** | **Braak stage** | **PMI** | **Cause of Death** |
| --- | --- | --- | --- | --- | --- |
| 1 | 69 | F | Stage 0 | 37h | Cardiac failure |
| 2 | 65 | F | Stage 0 | 5h | Pulmonary fibrosis |
| 3 | 73 | M | Stage 0 | 51h | Myocardial infarction |
| 4 | 39 | M | Stage 0 | 59h | Cancer |
| 5 | 85 | F | Stage 0 | 36h | Heart failure |
| 6 | 86 | F | Stage 3 | 67h | Cancer |
| 7 | 77 | F | Stage 3 | 59h | Lymphoma |
| 8 | 85 | F | Stage 3 | 95h | Cancer |
| 9 | 80 | M | Stage 3 | 48h | Cancer |
| 10 | 91 | M | Stage 3 | 23h | Septic shock |
| 11 | 91 | F | Stage 6 | 34h | Pneumonia |
| 12 | 74 | F | Stage 6 | 68h | Alzheimer's disease |
| 13 | 78 | M | Stage 6 | 44h | Alzheimer's disease |
| 14 | 81 | F | Stage 6 | 41h | Alzheimer's disease |
| 15 | 71 | F | Stage 6 | 64h | Advanced dementia |

Synaptosomes were prepared as previously described^40,61^. In brief, 300-400 mg of frozen tissue was homogenised with 1 mL, ice-cold homogenisation buffer (25 mM HEPES (pH 7.5), 120 mM NaCl, 5 mM KCl, 1 mM MgCl2, and 2 mM CaCl2, dissolved in HPLC-grade water), using a 2 mL Dounce homogeniser (Cambridge Scientific, Cat. 40401). Then the homogenate was serially filtered through an 80 µm nylon filter (Millipore, Cat. NY8002500; 25 mm filter holder: PALL, Cat. 4320) to remove tissue debris, followed by a 5 µm filter (Millipore, Cat. SMWP04700), to remove the organelles and the nuclei. The product was collected in a 1.5 ml Lo-bind Eppendorf and centrifuged at 1000 g for 5 minutes at 4□, the supernatant was removed, and the pellet was reconstituted with 100 uL homogenisation buffer.

### 2.2 SynPull imaging

SiMPull coverslips were prepared as previously described^36^. In brief, glass coverslips were first cleaned and passivated with polyethylene-glycol and coated with biotinylated polyclonal anti-neurexin 1 (NRXN1) antibody (10 nm in TBS with 1ug/mL BSA for 15 minutes; abcam, Cat. ab222806) using biotin-neutravidin interactions. Then the synaptosomes were incubated on the surface overnight at 4□. The next day, synaptosomes were fixed and permeabilised, and then synaptic AT8-positive tau aggregates were stained using Alexa Fluor^™^ 647 NHS ester labelled anti-AT8 antibody (2 nm in TBS with 1ug/mL BSA for 30 minutes; Invitrogen, Cat. MN1020B). For the co-localisation analyses, additional antibodies against T181-positive tau (Abcam, Cat. ab236458), phosphatidylserine (Sigma-Aldrich, Cat. 16-256), CD47 (Invitrogen, Cat. PA5-80435), and synaptogyrin-3 (Abcam, Cat. ab302614) labelled with Alexa Fluor^™^ 488 were added with the anti-AT8 antibody.. Lastly, the synaptosome membranes were stained using CellMask^™^ plasma membrane stain (Thermo Fischer Scientific, Cat. C10045) for 10 minutes (1:6000 by volume in TBS), followed by four rounds of washes with TBS. Direct stochastic optical reconstruction microscopy (*d*STORM) imaging for super-resolution size and shape analyses^40^ along with diffraction-limited single-molecule imaging^38^ were performed on a purpose-built total internal reflection fluorescence (TIRF) microscope^62^.

### SynPull image analysis

Super-resolution images acquired using *d*STORM were analysed using the Aggregate Characterisation Tool (ACT)^63^ as described previously^40^. Briefly, image preprocessing involved thresholding, dilation, and erosion to segment single molecules. The positions of segmented molecules were determined using ThunderSTORM^64^, and super-resolved images were generated by superimposing Gaussian-blurred localisations with a precision of less than 20 nm. Synaptosomes were segmented from CellMask images using a custom pipeline, as described previously^40^. Prior to synaptosome imaging, TetraSpeck fluorescent beads (ThermoFisher Scientific, Cat: T7279, diluted 1:1000 in PBS) were imaged in the same field-of-view (FoV) across both the CellMask (ThermoFisher Scientific, Cat: C10045; 561 nm) and antibody (640 nm) channels. These reference images were used to compute an affine matrix, which was subsequently applied to the CellMask images to correct for aberration-induced channel misalignment.

The corrected CellMask images were thresholded, and regions with intensities exceeding the mean plus two standard deviations within the FoV were identified as objects of interest. Noise-induced small objects were removed by erosion and dilation operations. The remaining objects were classified as synaptosomes following size filtering as described previously^40,65,66^. A density-based clustering algorithm (DBSCAN) with a 75 nm epsilon distance and a minimum of two points was applied on the super-resolution localisations in the 640 nm channel to identify synaptic and extra-synaptic aggregates.

Diffraction-limited fluorescence microscopy images were acquired for multiple targets of interest, each visualised in a separate channel (488 and 638□nm). Individual fluorescent puncta were segmented in each channel, and their centroid coordinates were extracted using a previously established method^40^. Using custom MATLAB scripts (R2024b, MathWorks), pairwise Euclidean distances were computed between puncta in different channels. For each punctum, the nearest neighbour distance (NND) to puncta in another channel was calculated. A distance threshold of 50□nm was applied. A punctum was considered colocalized if its NND to a punctum in another channel was less than or equal to this threshold. For each FoVs, we quantified how many puncta met this criterion, for example, the number of 561 puncta near 638, and those simultaneously near both 638 and 488.

### Tissue homogenization

For the ELISA and SIMOA analyses, human brain samples were homogenised as previously described^40^. In brief, ∼400 mg of tissue samples were placed into 2 mL Eppendorf protein Lo-bind tubes prefilled with 1 mm zirconium beads (Scientific Labs, Cat. SLS1414) and mixed with 700 uL of homogenising buffer (10 mM Tris-HCl, 0.8 M NaCl, 1 mM EGTA, 10% sucrose, 0.1% sarkosyl, Pefabloc SC protease inhibitor, and PhosSTOP phosphatase inhibitor tablet; pH∼7.4) and the samples were mechanically homogenised on an electronic tissue homogeniser (VelociRuptor V2 Microtube Homogeniser, Scientific Labs, Cat. SLS1401) at 5 m/s for two cycles of 15s, with a 10 s gap in-between, followed by centrifugation at 21,000 g for 20 min at 4□. Then the supernatant was collected and stored in a protein LoBind Eppendorf at 4□, while the pellet was homogenized again by adding an additional 700 uL of buffer and using the same parameters. The supernatant from this step was mixed with the one from the previous step, aliquoted, and stored in a −80□ freezer.

### Statistical analyses and data modeling

The R Project Statistical Computing version 4.4.2 (2024-10-31) -- “Pile of Leaves” was used for statistical analyses and the graphs were generated in GraphPad Prism 7.0a for Mac OS X. Differences between groups were determined using 95% confidence intervals (CI)^67^, presented on **Table 2**. No novel code was generated during data analysis and the datasets generated and analysed during the current study are available from the corresponding author on reasonable request.

**Table 2.** Statistical analysis for the comparison of the samples from different Braak stages. 95% confidence intervals of group differences are calculated and a difference not containing 0 is considered a statistically meaningful (significant) difference.

| **95% confidence intervals** | **Stage 0 vs 3** | **Stage 3 vs 6** | **Stage 0 vs 6** |
| --- | --- | --- | --- |
| **Tau synaptic fraction** | 2.85, 17.05 | 0.87, 15.07 | 10.82, 25.02 |
| **Synaptic tau count** | -2.80, 19.20 | -6.30, 15.70 | 1.90, 20.90 |
| **Synaptic tau length** | -18.30, 90.84 | 4.64, 51.68 | 11.00, 117.88 |
| **Extra-synaptic tau count** | 18.80, 144.40 | 157.70, 283.30 | 239.30, 364.90 |
| **Extra-synaptic tau length** | -8.70, 2.77 | -2.90, 4.40 | -7.36, 2.94 |
| **Total T181** | 3.33, 11.39 | -2.34, 5.72 | 5.02, 13.08 |
| **T181 synaptosome fraction** | 6.31, 18.72 | -3.88, 8.54 | 8.64, 21.06 |
| **T181+AT8 / AT8+ synaptosomes** | 2.41, 20.13 | -3.11, 14.61 | 8.21, 25.82 |
| **T181+AT8 / All synaptosomes** | 0.83, 9.80 | 1.51, 10.47 | 6.83, 15.79 |
| **Total PS** | 32.52, 47.23 | -23.79, -9.09 | 16.09, 30.79 |
| **PS synaptosome fraction** | 31.53, 48.28 | -18.58, -1.82 | 21.33, 38.08 |
| **PS+AT8 / AT8+ synaptosomes** | 21.52, 45.69 | -17.27, 5.55 | 15.84, 39.64 |
| **PS+AT8 / All synaptosomes** | 8.47, 23.15 | -11.76, 2.92 | 4.05, 18.73 |
| **Total CD47** | -16.98, -10.70 | -5.35, 0.93 | -19.19, -12.91 |
| **CD47 synaptosome fraction** | -27.10, -8.65 | -12.06, 6.38 | -29.94, -11.49 |
| **CD47+AT8 / AT8+ synaptosomes** | -27.72, 3.95 | -21.90, 8.91 | -34.78, -1.98 |
| **CD47+AT8 / All synaptosomes** | -0.06, 8.04 | -6.76, 1.33 | -2.77, 5.32 |
| **Total SYNGR3** | 19.43, 31.23 | 2.31, 14.11 | 27.64, 39.44 |
| **SYNGR3 synaptosome fraction** | 19.06, 38.51 | -1.95, 17.50 | 26.83, 46.28 |
| **SYNGR3+AT8 / AT8+ synaptosomes** | 15.93, 50.92 | -17.94, 13.91 | 16.52, 46.29 |
| **SYNGR3+AT8 / All synaptosomes** | 1.91, 15.24 | -3.24, 10.09 | 5.33, 18.66 |

We fit a geometric decay to the aggregate length distribution using a maximum likelihood fitting technique, based on the balance of elongation and aggregate removal^48^. Based on the assumptions of this model, we only perform the fit over a range of aggregate lengths, specifically, aggregates with more than _man_ monomers and less than _max_ monomers. Here, we choose _man_ = 500 and _max_ = 1200, and convert the aggregate lengths to monomers by multiplying the length of aggregate in nanometres by 4, from the assumption of a double stranded aggregate with a beta-sheet separation of 0.5 nm^68^. However, as shown Cotton et al.,^48^, the results for the difference in relative removal rate between synaptic and extra-synaptic aggregates is insensitive to this conversion factor. Our results are thus robust to the exact conversion and similarly apply if the aggregates differ in structure from those seen in cryo-EM, resulting in a different number of monomers per length. As such, we use the following normalised distribution to generate the probability of an aggregate of a length I as

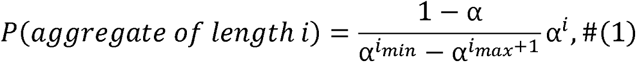

where α is the decay length and is to be determined. We determine α as the maximum likelihood for a distribution to generate all aggregates within length _man_ and _max_ . We perform this analysis for the intra- and extra-synaptic aggregate length distributions separately, across all Braak stages, to determine α_an ra_ and α_ax ra_. We define the normalized removal, r̃, as the removal rate/elongation rate. We determine the ratio of the synaptic and extra-synaptic relative removal using

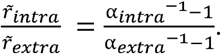

The quoted errors are for the 68^th^ percentile from the fitting of the decay lengths.

In order to quantify the effect of AT8-positive tau presence on the probability of encountering the other protein signal (phosphatidylserine, CD47, and synaptogyrin-3), we employed Bayesian inference. Since the data contain information on the presence and absence of a certain signal, the likelihood function is a Bernoulli distribution with unknown parameter *f*, which denotes the fraction of positive synaptosomes that one would expect to observe if infinite measurements are performed. Using a beta distribution as a prior, the resulting posterior probability of f is also beta distributed and simply given by the analytical expression:

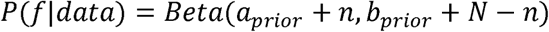

where *N* is the total number of synaptosomes, and *n* is the number of synaptosomes that contain the signal.

Furthermore, when comparing two distributions, such that of synaptosomes that contain the signal, and that of synaptosomes that contain the signal given they are also containing AT8-positive tau, we can evaluate the performance of the hypothesis that they were drawn from the same distribution, i.e. that *f _P_*_(*signal*+|*AT*8+)_ = *f*_p(_*_signal_*_+)_, with the hypothesis that they were not. In other words, we can work how likely it is that the presence of AT8-positive tau has an effect on the appearance of the signal. To do so, we compute the Bayes factor *B*, which is the probability of observing the data given that the two *f* parameters are the same, over the probability of observing the data given that the two *f* parameters are independent:

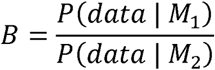

where *M_1_* and *M_2_* denote the different models, i.e. *M_1_* means *f _P_*_(*signal* +*AT*8 +)_ = *f _P_*_(_*_signal_*_+)_ and *M_1_* means *f _P_*_(*signal* +*AT*8+)_ and *f _P_*_(*signal* +)_ are independent. For easier interpretability we convert the Bayes factor to a probability that *M_1_* is correct, given by 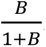. Thus a 50% probability corresponds to equal probabilities for *M_1_* being correct and *M_2_* being correct. Note that we analyse all biological repeats together, thus implicitly make the assumptions that *f* is constant across samples of the same disease stage. If in reality *f* differs between samples, the resulting posterior would be more spread out.

## 5. Declarations

### Ethical approval

Human post-mortem brain tissue was acquired from the Cambridge Brain Bank (Cambridge University Hospitals). The Cambridge Brain Bank is supported by the NIHR Cambridge Biomedical Research Centre (NIHR203312). We gratefully acknowledge the participation of all our patient and control volunteers.

### Availability of data and materials

Data collected during these experiments will be made available upon reasonable request from the corresponding author. No novel code was generated.

### Author contributions

**E.F.** Conception and design, data collection and statistical analysis, manuscript writing. **S.K.** Conception and design, data collection and analysis, manuscript preparation. **G.M.** and **M.W.C.** Data modelling. **A.Q.** Providing human brain samples, neuropathological characterisation. **M.G.S.** Conception and design, manuscript preparation. **D.K.** Conception and design, manuscript preparation, overall supervision of the project. **Acknowledgments:** We thank Dr. Kristy Halliday from the Cambridge Brain Bank for her assistance on human brain tissue acquisition.

