## Supplementary figures and images for "Early synaptic pathology is associated with small tau aggregates in Alzheimer’s disease"

### Supplemental Figure 1

#### Supplemental Figure 1

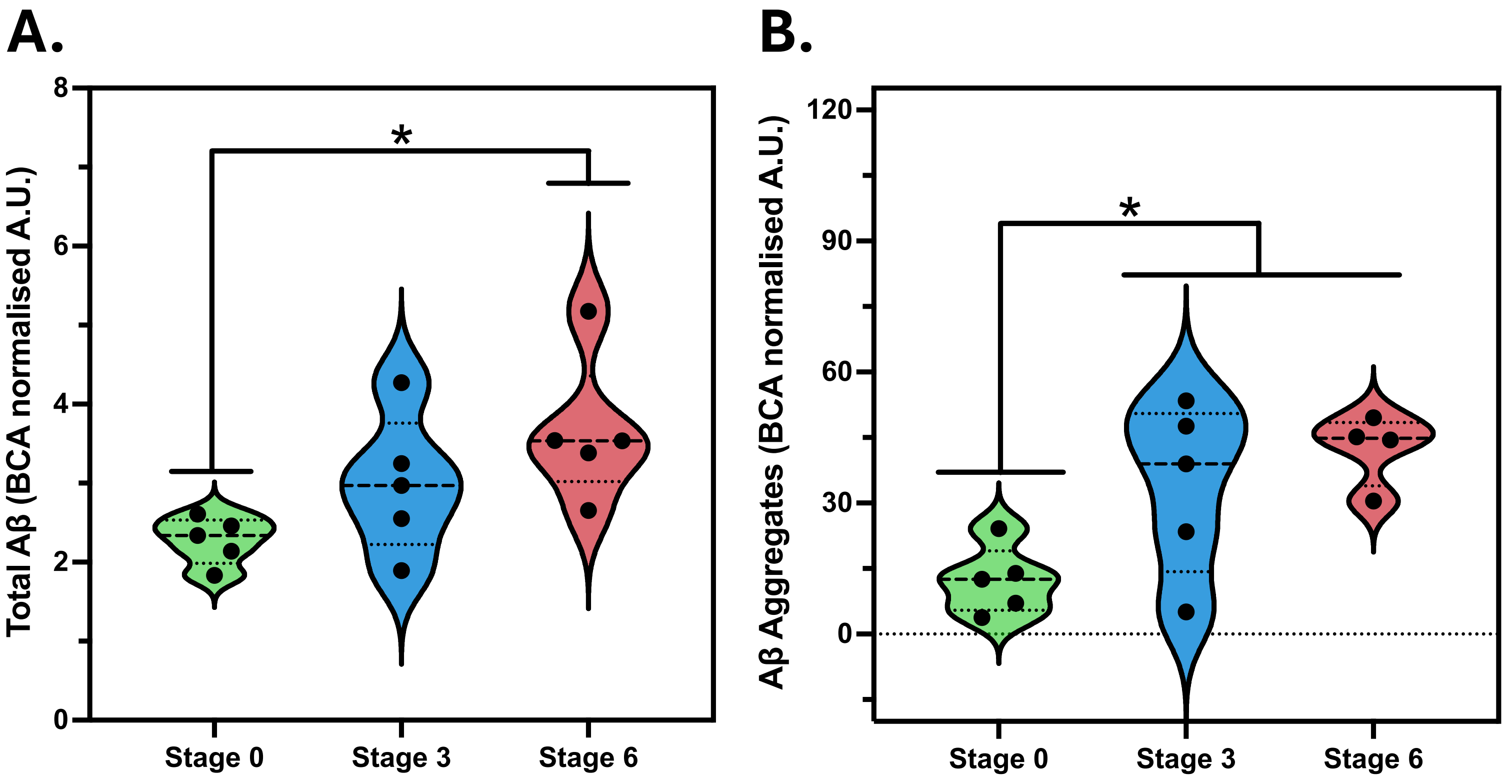
