## Supplemental text for "Early synaptic pathology is associated with small tau aggregates in Alzheimer’s disease"

28.07.2025

### Supplemental material:

While the current study focused on synaptic tau aggregates, it has been shown that oligomeric beta-amyloid (Aβ) can influence the synaptic accumulation and aggregation of tau. Thus, we have also quantified the total (monomer and aggregate) and aggregated diffusible Aβ concentrations in the same brain samples, using ELISA and purpose-built single-molecule array (SiMoA) assays, using the following methodology.

Total (monomeric and aggregated) soluble Aβ levels were determined using a commercial ELISA assay (Abcam, Cat: 289832), following the manufacturer's instructions. Meanwhile, the aggregate concentration in the same samples were determined using a SIMOA assay we have developed and validated previously[1,2]. In brief, brain homogenate samples were diluted to 10% in SIMOA lysate diluent A (Quanterix, Cat: 102908) and incubated with paramagnetic beads coupled to the monoclonal anti-Aβ antibody Aducanumab for 30 minutes at 37°C on a plate shaker. This antibody was chosen due to its binding profile for aggregates with a variety of shapes and sizes through different disease stages[1]. After washing, the beads were first incubated with biotinylated Aducanumab, forming an immunocomplex and then treated with streptavidin beta-galactosidase (SBG). Then the concentration of beads containing an immunocomplex as a ratio to overall beads were determined as a digital readout using the Quanterix SIMOA SR-X instrument.

While the total soluble Aβ levels did not show a difference between Braak stages 0 (control samples without neurofibrillary tangle pathology) and 3 ($CI_{95}$ = -0.33, 1.76) or 3 and 6 ($CI_{95}$ = -0.37, 1.71), there was significantly more soluble Aβ in the Braak stage 6 samples than stage 0 ($CI_{95}$ = 0.34, 2.43; **Supplemental Figure 1A**). On the other hand, Aβ aggregate levels showed a significant increase at stage 3 ($CI_{95}$ = 2.73, 40.10) but did not further increase by stage 6 ($CI_{95}$ = -11.13, 28.51; **Supplemental Figure 1B**). Together, these results show that an increase in soluble Aβ aggregate levels is an early event in AD and proceeds an increase in total (monomer and aggregate) Aβ levels.
